# Mortality attributable to seasonal influenza in Greece, 2013-2017: variation by type and age, and a possible harvesting effect

**DOI:** 10.1101/389411

**Authors:** Theodore Lytras, Katerina Pantavou, Elisavet Mouratidou, Sotirios Tsiodras

**Author notes:** Corresponding author: Theodore Lytras, MD, MPH Hellenic Centre for Disease Control and Prevention, Athens, Greece. Agrafon 3-5, 15123 Marousi, Athens, Greece.

## Abstract

**BACKGROUND:** Estimating the contribution of influenza to excess mortality in the population presents substantial methodological challenges. We combined environmental, epidemiological and laboratory surveillance data to estimate influenza-attributable mortality in Greece, over four seasons (2013-2014 to 2017-2018), specifically addressing the lag dimension and the confounding effect of temperature.

**METHODS:** Associations of influenza type-specific incidence proxies and of daily mean temperature with mortality were estimated with a distributed-lag non-linear model with 30 days of maximum lag, separately for each age group. Total and weekly deaths attributable to influenza and cold temperatures were calculated.

**RESULTS:** Overall influenza-attributable mortality was 23.6 deaths per 100,000 population (95%CI: 17.8, 29.2), and varied greatly between seasons, by influenza type and by age group, with the vast majority occurring in persons 65 years or older. Most deaths were attributable to A/H3N2, followed by type B influenza. During periods of A/H1N1 circulation, weekly attributable mortality to this subtype among older people increased rapidly in the first half, but then fell to zero and even negative, suggesting a mortality displacement (harvesting) effect. Mortality attributable to non-optimum temperatures was much higher than that attributable to influenza.

**CONCLUSIONS:** Studies of influenza-attributable mortality need to take distributed-lag effects into account, stratify by age group and adjust for circulating influenza types and daily mean temperatures, in order to produce reliable estimates. Our approach is useful and readily applicable in the context of influenza surveillance.

## Introduction

Seasonal influenza is a significant public health problem worldwide. A recent modelling study estimated that 291,000 – 646,000 annual deaths globally are attributable to influenza, most of those in older adults [1]. The impact of influenza on mortality can vary widely each year and across different populations, due to factors such as the types of circulating viruses, the age structure of the population, its level of health and access to healthcare, and its degree of immunity due to vaccination or previous influenza epidemics.

There is intense interest in obtaining accurate and timely estimates of influenza-attributable mortality across all age groups, in order to inform public health policy and evaluate the effectiveness of preventive interventions. Numerous studies have attempted to assess influenza-attributable mortality, mostly in developed countries, and with very heterogeneous results [2]. Differences in study methodology are important contributors to this heterogeneity, besides actual host- and pathogen-related factors [2].

The variety in the employed methods is a reflection of the many challenges involved in accurately estimating the part of population mortality that is attributable to influenza. First, the types of circulating influenza strains need to be taken into account in any statistical model, as not all types may have the same effects on all age groups [3]. This highlights the need to have solid laboratory data to inform an influenza activity proxy, as models without such a proxy tend to produce higher mortality estimates [2]. Second, estimation needs to adjust for the confounding effect of ambient temperature, not only at the extremes but also at milder cold levels, which are known to account for substantially more temperature-related deaths than extreme cold [4]. Finally, the association between influenza circulation and mortality is characterized by a lag that cannot be assumed to be fixed; not all people die at the same time after falling ill with influenza, therefore the effect of a given level of influenza activity on mortality is likely to be distributed over a period of time. Such distributed-lag associations with mortality have been previously decribed for temperature [4] and air pollution [5], but are also likely for influenza as well. To our knowledge though, very few studies to date have attempted to account for the lag dimension in the association between influenza and mortality [6–9], despite the availability of powerful relevant methods [10].

The objective of this study was to examine the mortality attributable to influenza over four winter seasons in Greece, and across different age groups, while addressing the methodological issues outlined above. For this purpose we combined epidemiological and virological data routinely collected as part of public health surveillance, with temperature data covering the entire country. A secondary objective was to compare the mortality attributable to cold temperatures with that attributable to influenza, in order to assess which one is the primary driver of excess winter deaths [11,12].

## Methods

### Data sources

The study encompassed the entire population of Greece, a developed country with a temperate climate and a population of slightly less than 11 million. Individual notifications of all-cause deaths from ISO week 22/2013 (May 2013) to week 41/2017 (October 2017) were available and obtained in December 2017 from the Hellenic Ministry of the Interior. In this dataset 99.3% of all deaths were registered within one week of their occurrence, thus there is practically no bias due to reporting delay. Deaths coded as violent (0.5% of the total) were excluded. We summed the notifications by age to create daily counts of deaths for three groups: all ages, 65 years or older, and 15-64 years. In the 0-14 years age group the data were too sparse (median 2 deaths/day, range 0-19) and were not analyzed further. Population denominators by age group were also obtained from the Hellenic Statistical Authority.

Average daily temperatures for the same period recorded at seven weather stations in Greece were downloaded from the website of the National Centers for Environmental Information (NCEI) of the National Oceanic and Atmospheric Administration (NOAA) [13]. A population-weighted daily mean over all stations was used as the countrywide temperature of that day.

Weekly Influenza-Like Illness (ILI) rates for the same period were obtained from the Hellenic Centre for Disease Control and Prevention (HCDCP). These rates are recorded by a sentinel surveillance system operated by the HCDCP, based on a countrywide geographically representative network of primary care physicians who voluntarily report every week their total number of patient consultations and the number of patients with ILI. As primary care physicians in Greece do not have a defined catchment area, the ILI rate denominator is total consultations rather than population. Aggregate laboratory data were also obtained by the two National Influenza Reference Laboratories operating in Greece (Hellenic Pasteur Institute, and Microbiology Laboratory, Aristotle University of Thessaloniki School of Medicine), namely the weekly number of all respiratory specimens tested by Real-Time PCR (RT-PCR) for influenza, and the number of specimens positive for A/H1N1, A/H3N2 and type B influenza. These two laboratories perform the bulk of influenza RT-PCR testing for surveillance purposes in Greece.

To create incidence proxies for type-specific influenza activity, we multiplied the weekly ILI rates by the proportion of specimens testing positive to each of the three major influenza strains, as previously suggested in the literature [14]. Under certain assumptions, these proxies are approximately linearly related to the weekly type-specific incidence of influenza in the population [15]. Each weekly time-series of influenza incidence proxies was converted to a daily time-series by assigning the weekly values to all days of the each week and applying a cubic smoothing spline with one degree of freedom per week, with the smoothing parameter chosen using the Generalized Cross-Validation (GCV) criterion. This method introduces minimal smoothing to the daily time series, essentially only removing the jags between adjacent weeks (Supplementary Figure 1).

### Statistical analysis

The association between daily death counts for each age group, average daily temperature and the three influenza incidence proxies was modelled using distributed-lag non-linear models (DLNM) of quasi-Poisson family and with log link [10]. In this type of models, both the “conventional” exposure-response and the additional lag-response association are simultaneously assessed using a pair of functions known as the cross-basis functions, which transform the exposure variable to a cross-basis matrix. The regression coefficients for this matrix, expressed as a Relative Risk (RR), then describe the relationship of the exposure to the outcome across both dimensions [10]. The general shape of both the exposure-response and the lag-response association needs to be described by specifying appropriate cross-basis functions, which can be non-linear. The maximum lag also needs to be specified.

The exposure-response curve for temperature was modelled with a quadratic B-spline with three internal knots placed at the 10th, 75th and 90th percentile of the temperatures distribution, similar to previous studies [4]. For the three influenza incidence proxies a linear relationship with mortality was specified; this implies a fixed case-fatality ratio per influenza subtype, i.e. that mortality is proportional to the incidence of infection with a particular subtype. We modelled the lag-response association for all four variables using a natural cubic spline with three internal knots placed at equally spaced values on the log scale [4]. The maximum lag was set to 30 days, in order to capture the delayed effects of both influenza and cold temperatures, but also in order to assess potential mortality displacement (harvesting) over this period, i.e. detect deaths that would occur anyway and were only brought forward in time by the exposure, rather than being “genuine” excess deaths that would not have occurred without the exposure [16]. The assumptions regarding the lag-response shape and the maximum lag were tested in a sensitivity analysis (Supplementary Figures 3-5).

Besides the four cross-basis matrices for temperature and influenza, we included in the regression models a linear term for trend, an indicator variable for day of the week, and a periodic cubic B-spline with three equidistant knots for day of the year (1-366) to model seasonality; the latter allows a more flexibly-shaped seasonal pattern of mortality compared to a classical Serfling-like sinusoidal term. One such model was fit on the daily death counts for every age group (all ages, 15-64, 65 or older).

For every day in the series, the number of deaths attributable to each influenza subtype and to cold temperatures was calculated using a previously described method for DLNMs [17]. Specifically, the attributable risk at each day was treated as being caused by multiple lagged effects of the exposure in the previous days up to the maximum lag back (“backward” attributable risk). The total attributable number of deaths is then given by summing the contributions of each day. Empirical 95% Confidence Intervals are calculated using Monte Carlo simulations, assuming a multivariate normal distribution of the cross-basis regression coefficients [17]. We used 50,000 replicates for that purpose, to ensure precision. We calculated influenza- and temperature-attributable mortality for each of the four winter seasons available in our dataset (defined as week 40 of each year to week 20 of next year, from 2013-14 to 2016-17), and also for every week in the study period to create an epidemic curve.

All calculations were performed with the R software environment, version 3.4.3 [18], and with package “dlnm” for R, version 2.3.2 [19].

## Results

From May 2013 to October 2017 a total of 518,688 deaths were recorded in our dataset, of whom 314,554 (60.6%) occurred during the four winter seasons. Most of these were in people aged 65 and older, 271,510 vs 40,639 in people aged 15-64. The four seasons were heterogeneous with respect to influenza activity (Figure 1). During season 2013-14, influenza A/H1N1 was dominant with some additional A/H3N2 activity; season 2014-15 was biphasic, with an early wave of A/H3N2 and an equally large second wave of type B influenza; season 2014-15 was completely dominated by A/ H1N1 with little late-season type B and virtually no A/H3N2; and season 2015-16 was dominated by A/H3N2 with a smaller late wave of type B. Average daily temperature showed a clear seasonal pattern, with some winters being warmer (e.g. 2013-14) and others having more colder days (such as 2016-17). Mortality in people aged 65 and over showed peaks during the winters of 2014-15 and 2016-17, coincidental with influenza A/H3N2 activity, while a more subtle peak in mortality for people aged 15-64 was observed in the winter of 2015-16, when A/H1N1 activity was high (Figure 1).

**Figure 1:**
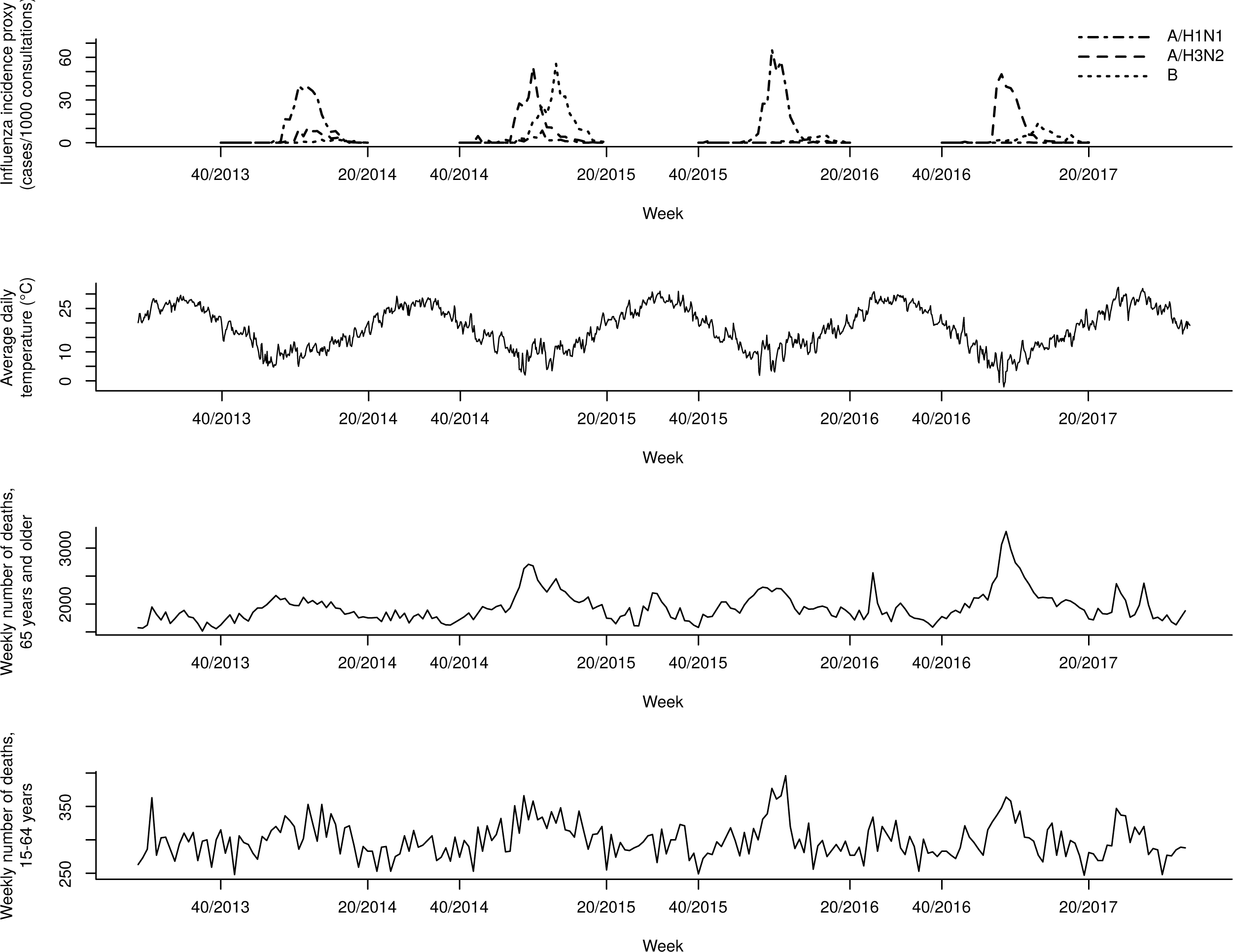
Influenza activity, average daily temperature and weekly all-cause mortality by age group, Greece, May 2013 to October 2017.

The overall association (across all lags up to the maximum of 30 days) of temperature with mortality, as provided by the fitted DLNMs, is illustrated in Figure 2. It was found to be slightly attenuated for those aged 15-64, although the difference was non-significant. The temperature of minimum mortality was 25°C, for both age groups. With respect to the three influenza subtypes, the overall RR is a linear function of the respective incidence proxies and it is illustrated on Table 1 for an indicative value of 30 cases per 1000 consultations, approximately corresponding to a medium intensity of influenza circulation. Type B influenza was associated with mortality in both age groups, but A/H3N2 appeared to affect only those aged 65 or older, (RR=1.14, 95% CI 1.09 to 1.19), while A/H1N1 affected only people aged 15-64 (RR=1.09, 95% CI 1.03 to 1.15).

**Table 1:** Overall association (Relative Risk) of influenza subtypes with all-cause mortality, across all lags, by age group, for an incidence proxy of 30 cases per 1000 patient consultations.

|  | A/H1N1 | A/H3N2 | B |
| --- | --- | --- | --- |
| All ages | 1.02 (1.00 - 1.04) | 1.12 (1.08 - 1.16) | 1.08 (1.05 - 1.11) |
| 65 years or older | 1.01 (0.99 - 1.03) | 1.14 (1.09 - 1.19) | 1.08 (1.05 - 1.11) |
| 15-64 years | 1.09 (1.03 - 1.15) | 1.00 (0.91 - 1.11) | 1.13 (1.05 - 1.21) |

**Figure 2:**
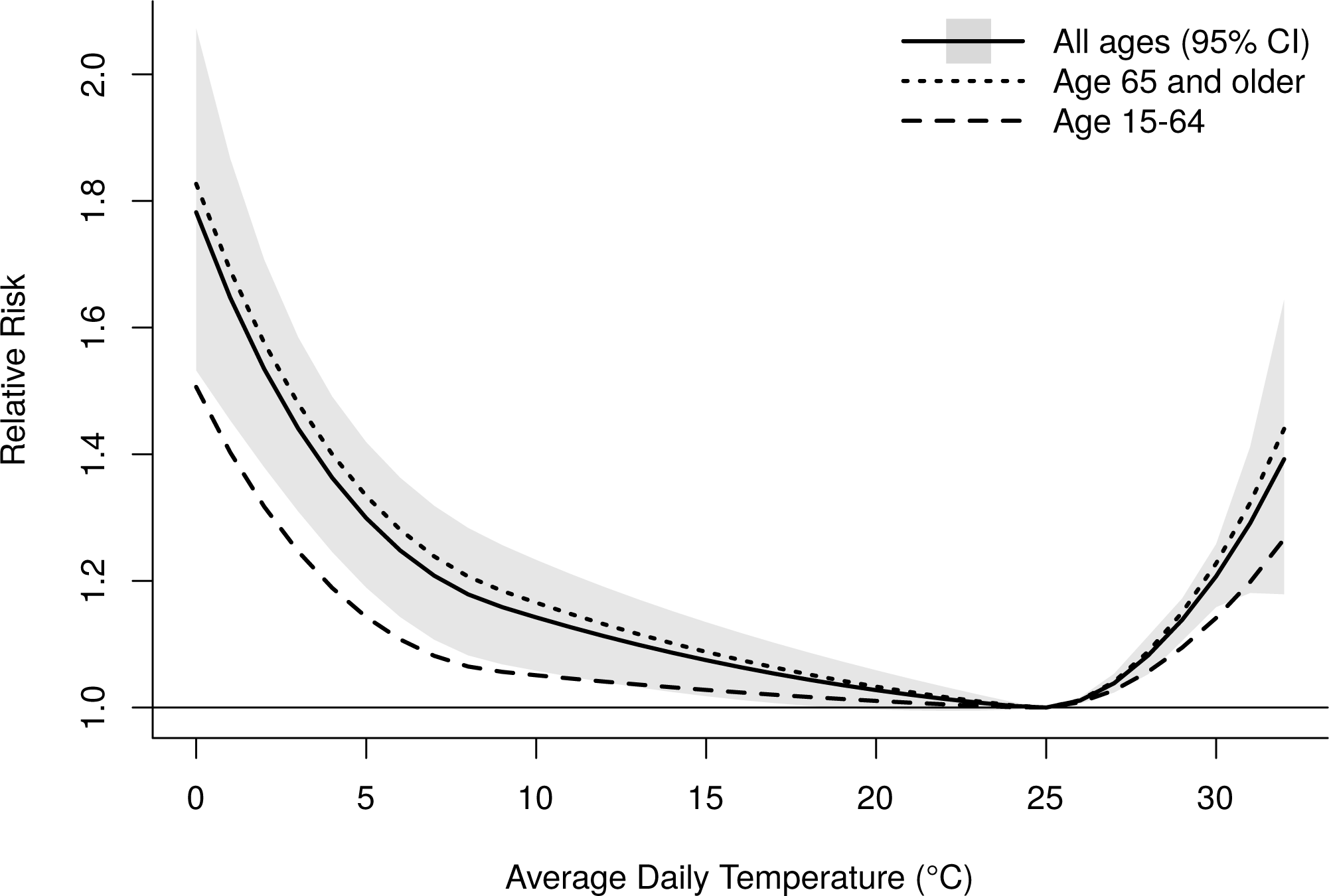
Overall association (Relative Risk) of temperature with all-cause mortality, across all lags and by age group.

Table 2 translates the above associations into attributable mortality per 100,000 population, overall and by season. Across all ages, influenza was associated with an annual mean of 2,559 excess deaths, or 23.6 per 100,000 population (95%CI 17.8, 29.2). The majority of these deaths occurred in people aged 65 and older (89.5%, or 2290 deaths per year), and the remainder in people aged 15-64 years. In comparison, cold temperatures (lower than the minimum mortality temperature) were associated with 74.7 excess deaths per 100,000 population (95%CI 35.3, 111.7), and accounted for the larger part of the wintertime seasonal excess mortality (Figure 3). Most of the influenza-attributable mortality was accounted for by subtype A/H3N2 affecting older people, with type B also making a substantial contribution. In contrast, even in seasons with high A/H1N1 activity, influenza A/H1N1 was not associated with a net excess of deaths among people 65 years and older (Table 2).

**Table 2:**
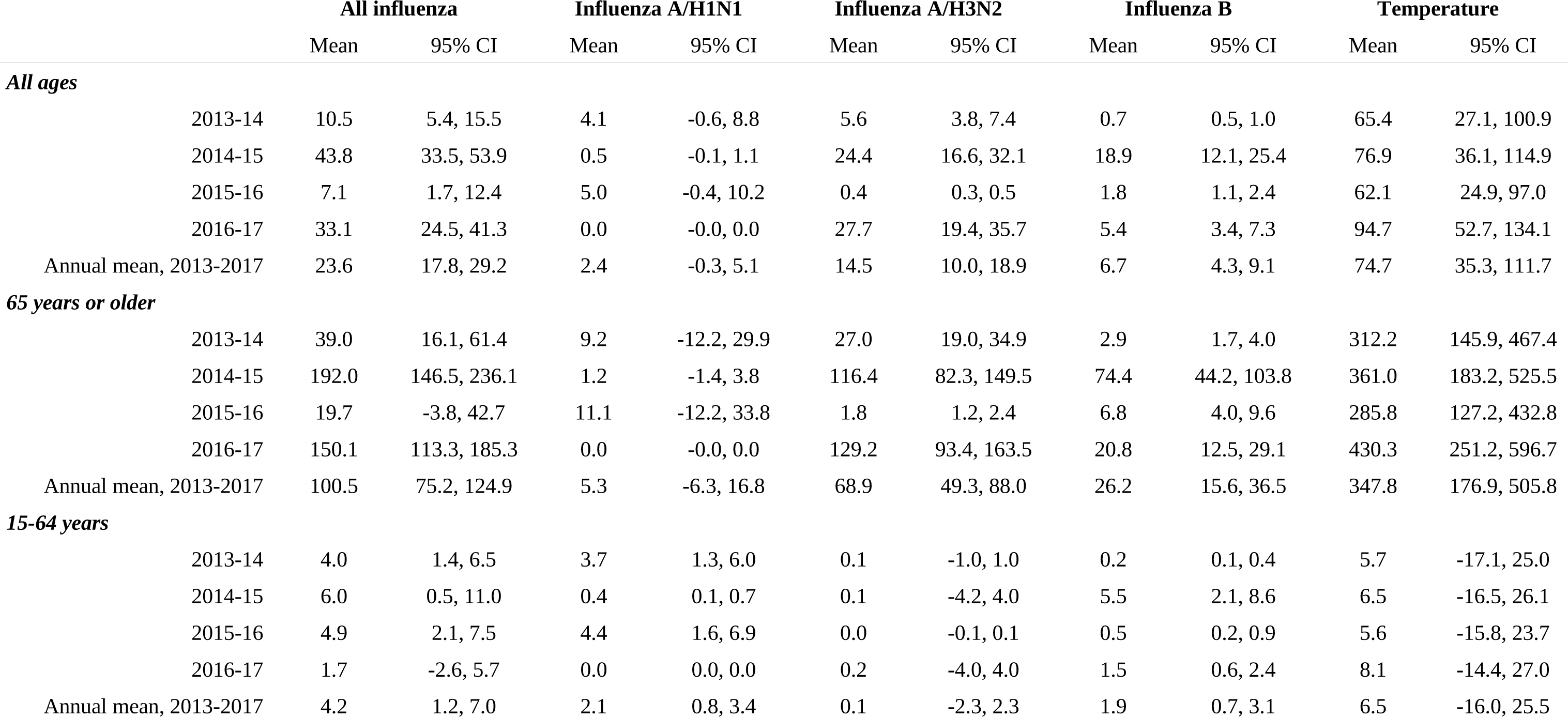
Mortality (per 100,000 population) attributable to influenza and temperature, by age group and season.

**Figure 3:**
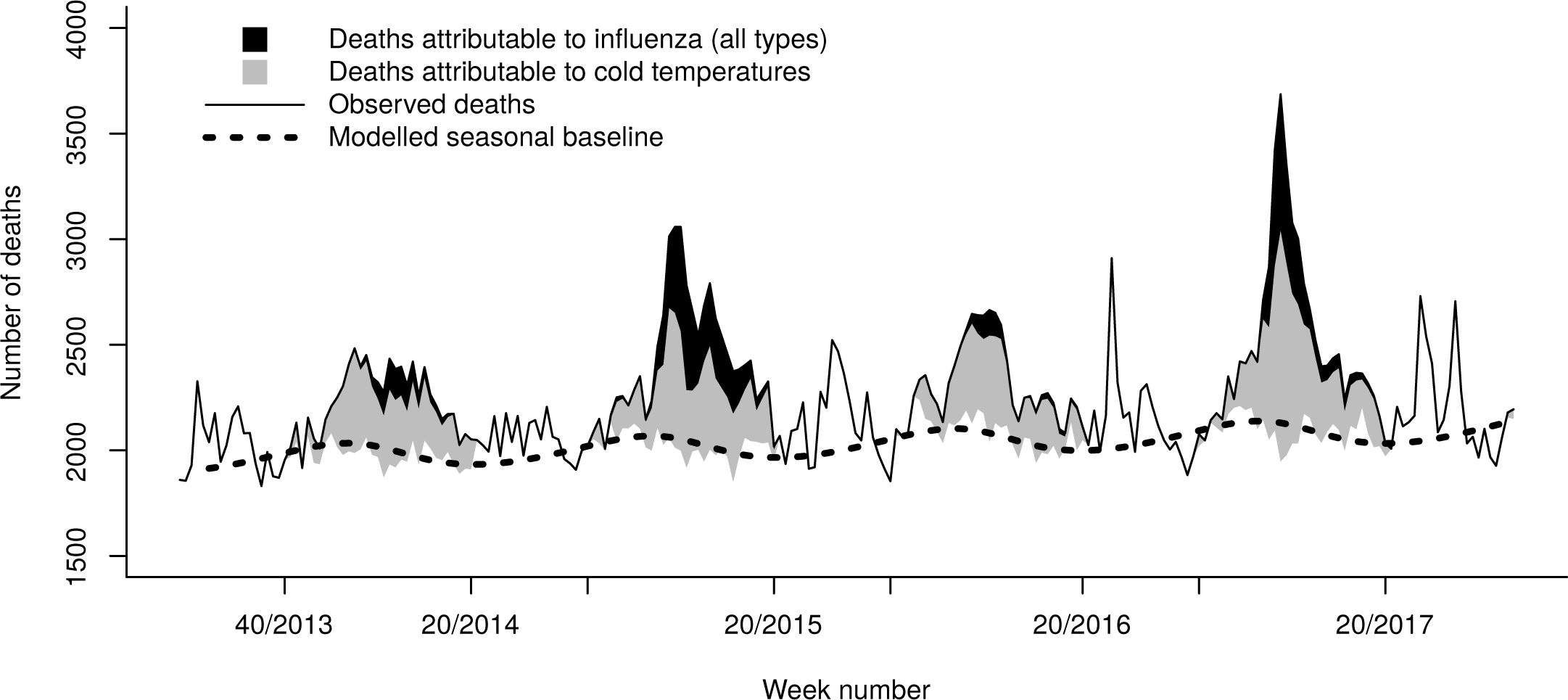
Weekly observed all-cause deaths, deaths attributable to influenza and to cold temperatures, Greece, May 2013 to October 2017.

However, when plotting weekly influenza-attributable mortality by subtype for the entire study period (Figure 4), an interesting pattern emerged: A/H1N1 activity appeared to be associated with a substantial excess of influenza-attributable deaths for several weeks, but towards the end of the seasonal epidemic wave the number of associated deaths fell to zero and even negative (for season 2015-16). Even though confidence bands were wide, this is suggestive of a possible mortality displacement (harvesting) effect for influenza A/H1N1. Stratifying weekly A/H1N1-attributable mortality by age group shows this pattern only among those 65 years or older, and not among people aged 15-64 (Figure 5). In terms of mortality per 100,000 population of age 65 and older, in 2013-14 there was an excess of 11.2 deaths (95% CI −1.7 to 23.9) for the first 9 weeks (week 01/2014 to 09/2014) followed by a deficit of −2.1 deaths (95% CI −12.5 to 8.2) over the next 9 weeks. Similarly, in 2015-16 there was an excess of 14.7 deaths per 100,000 (95% CI 0.1 to 29) also over nine weeks (51/2015 to 06/2016), followed by a deficit of −3.6 deaths (95% CI −15.9 to 8.4).

**Figure 4:**
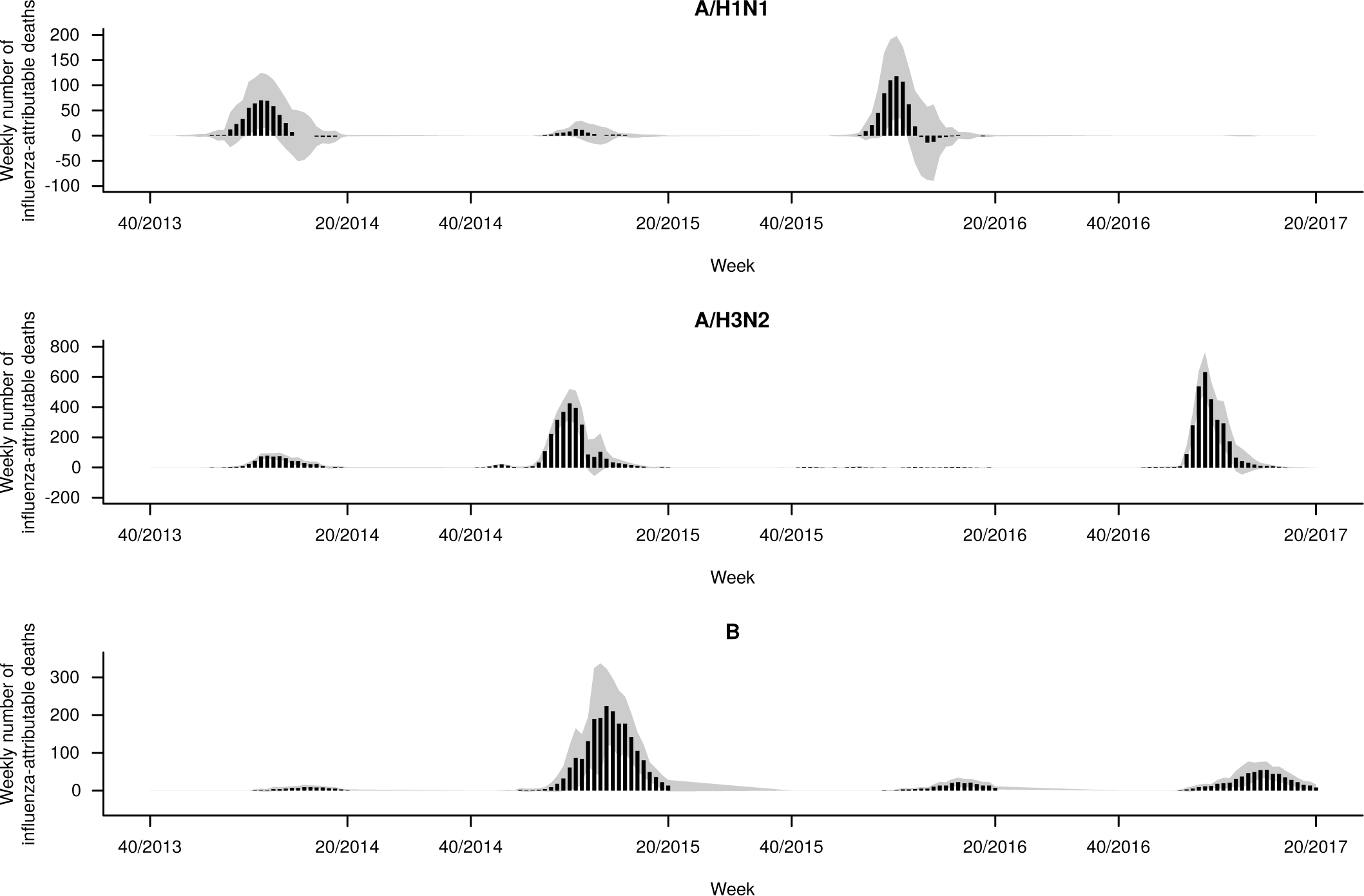
Epidemic curve of weekly influenza-attributable deaths for all ages, by influenza subtype. Four winter seasons in Greece, weeks 40/2013 to 20/2017.

**Figure 5:**
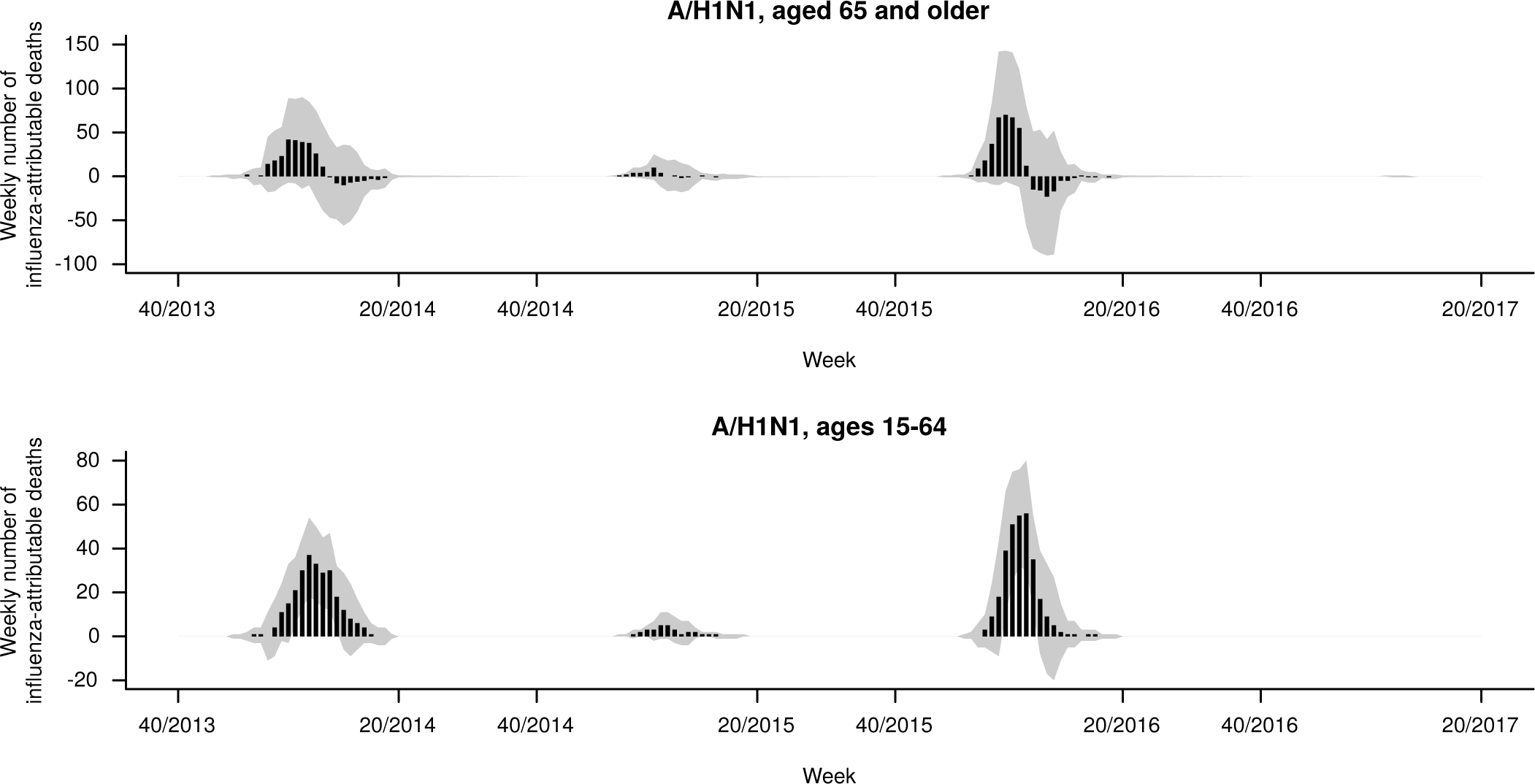
Epidemic curve of weekly deaths attributable to influenza A/H1N1, by age group. Four winter seasons in Greece, weeks 40/2013 to 20/2017.

Similar results were obtained in sensitivity analyses under alternative model specifications (Supplementary Figures 3–4). A longer maximum lag appeared to increase the number of deaths attributable to influenza A/H3N2 and augment the observed mortality displacement for A/H1N1, while a shorter maximum lag appeared to attenuate it. In addition, a model without the cross-basis matrix for temperature resulted in twice as many deaths attributable to influenza (all types) compared to the main model (4,552 vs 2,559 per year – Supplementary Figures 3 and 5).

## Discussion

Our results confirm that the mortality attributable to influenza varies greatly between seasons, and indicate that the types of circulating strains are a major determinant of this variation, with A/H3N2 being associated with many more deaths compared to A/H1N1 and type B influenza. As such, it is essential to individually account for each influenza type when modelling their association with mortality, using detailed laboratory data, rather than consider all influenza together in a single incidence proxy. We have also confirmed that the majority of influenza-attributable deaths occur among persons 65 years and older, and shown that each influenza type has different effects on mortality among different age groups, with A/H3N2 affecting older adults, and A/H1N1 younger persons. Therefore, in order to fully explore the effects of influenza on population mortality, stratification by age group should always be performed. Although it is difficult to compare studies due to different methodologies, our results from Greece are overall very similar to analogous estimates from the United States, with respect to both the total mortality burden attributable to influenza and the relative contribution of each influenza type [14]. They are also within the range reported in a recent systematic review of studies examining influenza-attributable mortality [2].

The addition of the lag dimension in the association between influenza and all-cause mortality offers a novel and important insight. To our knowledge this is the first time a short-term harvesting effect is suggested for influenza, for a particular type and in a particular age group. Such an effect could not have been detected by previous studies that did not employ distributed-lag modelling, and only considered fixed lags selected according to goodness of model fit [2]. One prior study from Sweden did observe that high winter mortality, including deaths coded as influenza, led to fewer heat-related deaths over the following summer [20]. Our study, however, is the first to directly associate with influenza A/H1N1 an excess of deaths among people 65 years or older, over several weeks, followed by a period with no excess or even a possible mortality deficit for several more weeks within the same season. Due to low precision, this finding will need to be confirmed in further studies applying similar distributed-lag models in bigger datasets from larger populations. If confirmed, it would suggest that among the elderly influenza A/H1N1 affects those with the most vulnerable health, many of whom would have died in the short-to-medium term from other causes independent of influenza. In contrast, A/H3N2 and type B appear to be associated with a “genuine” excess of influenza-related deaths, among older people who would not otherwise have died in the same year. This has important implications for correctly assessing the effect of circulating influenza types on population mortality, as well as the potential benefit of targeted vaccination campaigns.

There is an ongoing debate about whether the wintertime seasonal mortality excess is primarily driven by seasonal influenza [11,21] or cold temperatures [12,22]. In our study we found that mortality attributable to cold temperatures was several times higher than that attributable to influenza, with temperature explaining most of the within-year seasonal variation in mortality (Supplementary Figure 5). The strong confounding effect of temperature on the association between influenza and mortality is highlighted by our sensitivity analysis, which found a doubling of influenza-attributable deaths when not adjusting for temperature. As a result, to minimize residual confounding we specified the same maximum lag and lag-response specification for both influenza and temperature, and used daily counts of death and daily temperatures rather than weekly averages. On the other hand, long-term temperature trends are themselves highly seasonal (Figure 1), therefore it has been argued that distributed-lag models with long lags may capture part of this seasonality and produce larger effects of cold temperatures due to collinearity [21]. However, in our analyses we did not detect substantial collinearity (Variance Inflation Factors for all coefficients of interest was under 1.7), while in sensitivity analyses models with shorter or longer lag produced similar attributable mortality estimates for both influenza and temperature (Supplementary Table 3). Although it is true (and our results also indicate) that the magnitude of the overall winter excess mortality does correlate with the dominant influenza subtype for each season [11], influenza-attributable deaths appear to be a minority among all winter deaths. The majority is attributable to cold temperatures, independent of influenza, and that part is relatively constant each season.

Our proposed approach uses routine epidemiological and laboratory surveillance data, provides detailed weekly estimates of influenza-attributable mortality, and can be readily implemented with open-source software. As a result, it is particularly useful and applicable in an influenza surveillance context [23]. Further research and evaluation in settings other than Greece is warranted in this regard, as well as a comparison with other methods to estimate influenza-attributable mortality that are presently used in Europe [24].

Besides the advanced DLNM statistical methodology, a strength of the current study is the integration of comprehensive epidemiological, laboratory and weather data in a single detailed analytical model, offering more reliable estimates of influenza-attributable mortality. Thus we could account for individual influenza types, closely adjust for daily mean temperatures, flexibly model seasonality using periodic splines, and allow for distributed-lag effects. We also used daily death counts as the outcome, rather than the usual weekly counts, in order to minimize residual confounding.

On the other hand, the study has important limitations. With only four seasons of available mortality data on a relatively small population, statistical power to detect associations was suboptimal particularly for the younger age groups. Especially in the highly interesting 0-4 year group, deaths were so sparse that no informative conclusions could be drawn. In addition, we had only one season with significant influenza B circulation (2014-15, entirely of B/Yamagata lineage), which might not necessarily be representative of the actual effects on mortality of that influenza type, or of the B/Victoria lineage. We could only study all-cause deaths, as no comprehensive coded cause-specific mortality data are available in Greece; for example, it would have been very useful to examine cardiac or Pneumonia & Influenza (P&I) deaths, but unfortunately this was not possible. We also had no available laboratory surveillance data on Respiratory Syncytial Virus (RSV) infection; RSV is known to account for a substantial part of winter excess deaths [8,9], and might co-circulate with influenza [25], thereby confounding its association with mortality. Finally, we did not specifically consider the effect of seasonal influenza vaccination in the population; however, vaccination coverage in Greece is reportedly very low even among high risk groups [26], and is also unlikely to vary significantly between seasons.

In conclusion, we have analyzed the mortality burden attributable to influenza in Greece, and shown it to be significant especially among older people, as well as highly variable depending on the circulating influenza types. We have also shown how the DLNM framework can be used to explore the lag dimension, and identify potential mortality displacement in the association between influenza and mortality. More studies with similar methodology in other countries are warranted, to confirm these results, further study the observed harvesting effect, and evaluate the utility of this approach in the context of routine influenza surveillance. A final takeaway from this study: if the majority of influenza-attributable deaths occur in the age group (65 years and older) that has the lowest immune response to the seasonal influenza vaccine [27], and by the subtype (A/H3N2) that the vaccine usually offers the least protection against [28], then improved and more effective influenza vaccines are urgently needed to protect the population, with a view to the ultimate goal of a “universal” influenza vaccine [29].

